# Biochemical analysis of human eIF4E-DCP2 interaction: Implications for the relationship between translation initiation and decapping

**DOI:** 10.1101/2025.03.21.644491

**Authors:** Zachary F. Mandell, Jeff Coller

## Abstract

All eukaryotic mRNAs bear a 7-methylguanosine cap on their 5’ end. The 5’ cap enables mRNA translation by binding directly to eIF4E; which further recruits other factors and the 40S ribosome. Additionally, the 5’ cap maintains transcript stability; removal of the cap by the enzyme Dcp2 is necessary to degrade the mRNA. An *a priori* conclusion, therefore, has been that cap binding by eIF4E and DCP2 are antithetical to each other as both need access to the same substrate, i.e. the 5’ cap. In this study, we purified native full-length human eIF4E and Dcp2 and utilize biophysical and biochemical approaches to examine the in vitro interplay between Dcp2 and eIF4E. We confirm that Dcp2 is sufficient to remove the 5’ cap. Moreover, we demonstrate that Dcp2 binds RNA with nanomolar affinity. We discovered that, unexpectedly, eIF4E does not interfere with Dcp2’s decapping function, contradicting previous mechanistic models. Moreover, eIF4E binding appears to increase the affinity of Dcp2 for RNA. Although limited to in vitro conditions, our findings warrant a reevaluation of the proposed relationship between these mRNA cap-binding proteins.

## INTRODUCTION

A distinguishing feature of eukaryotic mRNA is the 7-methylguanosine (m7G) cap added co-transcriptionally to the 5’ end [1]. The 5’ m7GpppN cap serves two critical functions; it promotes ribosome association by binding the translational initiation factor eIF4E [2], and it protects the mRNA from degradation by blocking the activity of the 5’-to-3’ exoribonuclease, Xrn1 which acts preferentially on a 5’ monophosphate [3–5].

The control of mRNA stability is well recognized to be intimately connected with mRNA translation [4–10]; mRNAs that are decoded efficiently are more stable, while mRNAs that decode inefficiently are less stable. The first step in eukaryotic mRNA decay is removal of the 3’ polyadenosine tail by the CCR4/NOT complex [5]. This in turn allows for decapping of the 5’ cap by the DCP2 decapping enzyme and then digestion of the mRNA’s body 5’ to 3’ by XRN1 [5]. Much work has investigated the nature of how mRNA translation impacts mRNA decay rate [5]. In budding yeast, a major determinate of deadenylation and decapping rate is codon optimality which disproportionally alters elongation rate, allowing probing and binding of the ribosome by the CCR4/NOT complex [5]. In human cells, CCR4/NOT appears to also probe and bind the ribosome; codon specificity is also a determinant [5,11]. Following these events, a decapping complex must assemble on or in proximity to the 5’ cap structure, thereby allowing the cap to be enzymatically cleaved by DCP2. Importantly, since the process of mRNA occurs co-translationally [5], the presence of translational initiation factors on the 5’ end has theoretically been seen as a barrier to the decapping reaction [10]. Thus, it has been proposed that DCP2 binding and catalytic activity are mutually exclusive with eIF4E binding, although this has not been rigorously established [2,18]. A previous study suggested that an enzymatic mRNA decapping activity in yeast is inhibited in vitro by the addition of eIF4E [10], although this work predates the discovery of Dcp2 and thus requires reexamination.

Herein, we purified full-length human Dcp2 and eIF4E. We confirmed as previously reported that Dcp2 operates specifically on m7G capped RNA, with a turnover rate 5-fold greater than that of the yeast Dcp2:Dcp1 heterodimer [14]. This activity may potentially be explained by a tight binding affinity between human Dcp2 and RNA, as we find that Dcp2 exhibits an affinity of ∼157 nM for m7G-capped RNA, which is approximately the same as the affinity of eIF4E for m7G-capped RNA [29,31]. Intriguingly, we find that Dcp2 binds ∼1.6-fold more tightly to its enzymatic product, 5’ monophosphate RNA. Moreover, we find that eIF4E does not inhibit Dcp2 binding in our hands in vitro. In fact, eIF4E occupancy increases the affinity of Dcp2 for the RNA by ∼2-fold. Lastly, we find that pre-incubation of the RNA with eIF4E did not significantly impact the rate of decapping by Dcp2. Together, our data provide a few new insights into the relationship between eIF4E and the human mRNA decapping enzyme that should be considered as we further elucidate the interplay between translation and mRNA decay.

## RESULTS

### Human Dcp2 specifically removes the RNA 5’ m7G cap in the absence of decapping coactivators

Much is known about the activity of the eukaryotic decapping enzyme Dcp2 and its associated coactivators [12,13]. Dcp2 consists of an N-terminal regulatory domain (RD), a catalytic domain (CD), and a C-terminal intrinsically disordered region (IDR) [13]. The length of the RD and the IDR are highly variable across species and the various functions of these features are not entirely understood [13]. The CD is composed of a NUDIX hydrolase, which is a ubiquitous family of nucleotide hydrolases that typically utilize a metal cation to catalyze the hydrolysis of nucleosides [13]. In yeast, the C-terminal IDR auto-inhibits the enzyme [14] and binds additional mRNA decay factors [14–17]. In yeast, Dcp2 requires decapping coactivators, such as Dcp1, to achieve decapping [14,18–22]. Importantly, Dcp2 functions distinctly between eukaryotes. First, the C-terminal IDR of human Dcp2 is approximately 400 amino acids shorter than that of the yeast Dcp2 C-terminal IDR [18]. Second, Dcp2 does not form a heterodimer with Dcp1 in metazoa, rather these two factors interact indirectly via the scaffold protein Edc4 [23–25]. Third, biochemical studies of human Dcp2 have shown it does not require coactivators to achieve an appreciable level of decapping *in vitro* [26–29].

In order to understand the in vitro characteristics of the human decapping enzyme, we purified full-length Dcp2 to >90% purity (S1A Fig). We first tested enzyme activity by measuring decapping of a 43-mer RNA oligo that was modified at the 5’ end with m7GpppAm and labelled at the 3’ end with a Cy5 tag (S1B Fig). Purified Dcp2 was incubated for 30 minutes at 37°C with the synthetic RNA substrate. The reaction conditions were similar to those reported previously [26,27,32–34]. Resolution of the RNA substrate by PAGE revealed that incubation with Dcp2 changed the synthetic RNAs mobility, resulting in the appearance of a faster migrating species. This new species was comparable in size to a 43-mer RNA oligo that was identical in sequence, yet 5’ triphosphate (Fig 1A, lanes 2 vs. 3). Removal of m7GDP from the capped RNA was verified by incubation of the RNA with both Dcp2 and Xrn1. Xrn1 is a 5’ to 3’ exoribonuclease that acts specifically on 5’ monophosphate RNA [3]. Incubating the synthetic RNA with both Dcp2 and Xrn1 resulted in the disappearance of the faster migrating RNA species but not the slower species. These data suggest that the faster migrating species corresponded to RNA oligo that had been decapped by Dcp2 (Fig 1A lanes 3 vs. 5). To further verify the activity of purified Dcp2, we incubated the capped RNA substrate in the absence of divalent cations, where we confirmed that Dcp2 requires the presence of divalent cations in the reaction buffer to catalyze decapping (Fig 1B). Intriguingly, we also noted that Mg^2+^ coordinated the reaction more efficiently than Mn^2+^ (Fig. 1B, 73% vs. 59% decapped), which is inconsistent with previous reports using purified human Dcp2 [32]. Lastly, we confirmed that Dcp2-mediated catalysis is specific for the m7G cap. More specifically, we found that incubation of Dcp2 with RNA bearing either an ApppAm cap or 5’ triphosphate did not result in any alteration to the RNA (Fig 1C). Together, these results are consistent with previous observations that human Dcp2 decaps RNA in the absence of known decapping coactivators in vitro [26,27,32–34].

**Figure 1.**
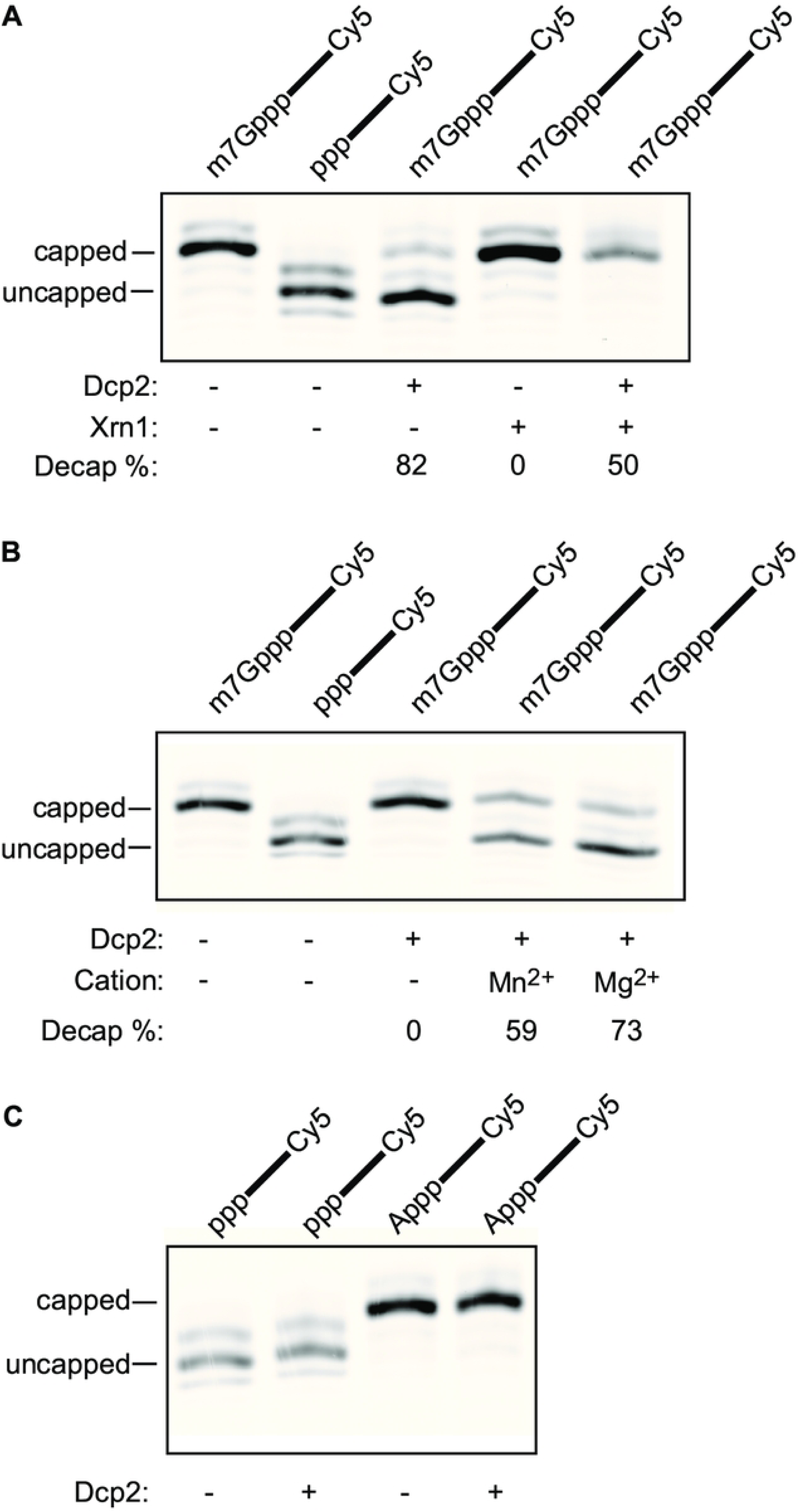
Human Dcp2 de-caps m7G-capped RNA. A. Decapping and decay reactions using RNA oligos containing the 5’ moieties that are indicated above each lane. Experiments were performed in the absence (-), or presence of Dcp2, and/or yeast Xrn1 as specified under each lane. The decapping % for the data shown here is shown underneath each lane. The capped and uncapped RNA species are indicated by the horizontal bars to the right of the gel slice. B. Decapping reactions using RNA oligos containing the 5’ moiety indicated above each lane. Experiments were performed in the absence or presence of Dcp2, Mg^2+^, and/or Mn^2+^, as indicated under each lane. The decapping % for the data shown here is shown underneath each lane. The capped and uncapped RNA species are indicated by the horizontal bars to the right of the gel slice. C. Decapping reactions using RNA oligo containing the 5’ moiety indicated above each lane. Experiments were performed in the absence or presence of of Dcp2 as indicated under each lane. The capped and uncapped RNA species are indicated by the horizontal bars to the right of the gel slice.

### Human Dcp2 displays affinity for RNA

Human Dcp2 has been reported to bind to both capped and 5’ triphosphate RNA [23,29]. To determine the affinity of Dcp2 for RNA, we first conducted a gel-shift assay using m7G-capped, 3’ unlabeled RNA in the presence of varying concentrations of Dcp2 (Fig 2A). To ensure that Dcp2 did not de-cap the RNA during the reaction, we omitted a source of Mg^2+^ (see Fig 1B). We found that incubation of the RNA with Dcp2 led to multiple distinct bound species, indicating that Dcp2 binds to RNA in a multivalent stoichiometry. To measure the affinity of these interactions, we utilized MicroScale Thermophoresis (MST) [35]. While the K_D_ fit is appropriate to model binding interactions that follow a 1:1 stoichiometry, the Hill fit is better suited to model multivalent interactions [36]. Considering the multivalency between Dcp2 and the RNA (see Fig. 2A), we chose to model our MST data using the Hill fit. The Hill fit yields two binding parameters, the EC_50,_ which is a measure of binding affinity, and the Hill coefficient [36]. Hill coefficients greater than 1 indicate a positive cooperativity [35], or a promiscuity of binding [37]. Quantitation of binding affinity is contingent on experimental conditions [38]. Thus we refer to all EC_50_ values in this report as the observed EC_50_, or the EC_50,obs_. Using this method, we found that the EC_50,obs_ between Dcp2 and m7G-capped RNA is 157±26 nM (Fig 2B). In addition, the Hill coefficient of this interaction was found to be 1.56±0.49, indicating that Dcp2 may bind to RNA nonspecifically, as others have suggested [39].

**Figure 2.**
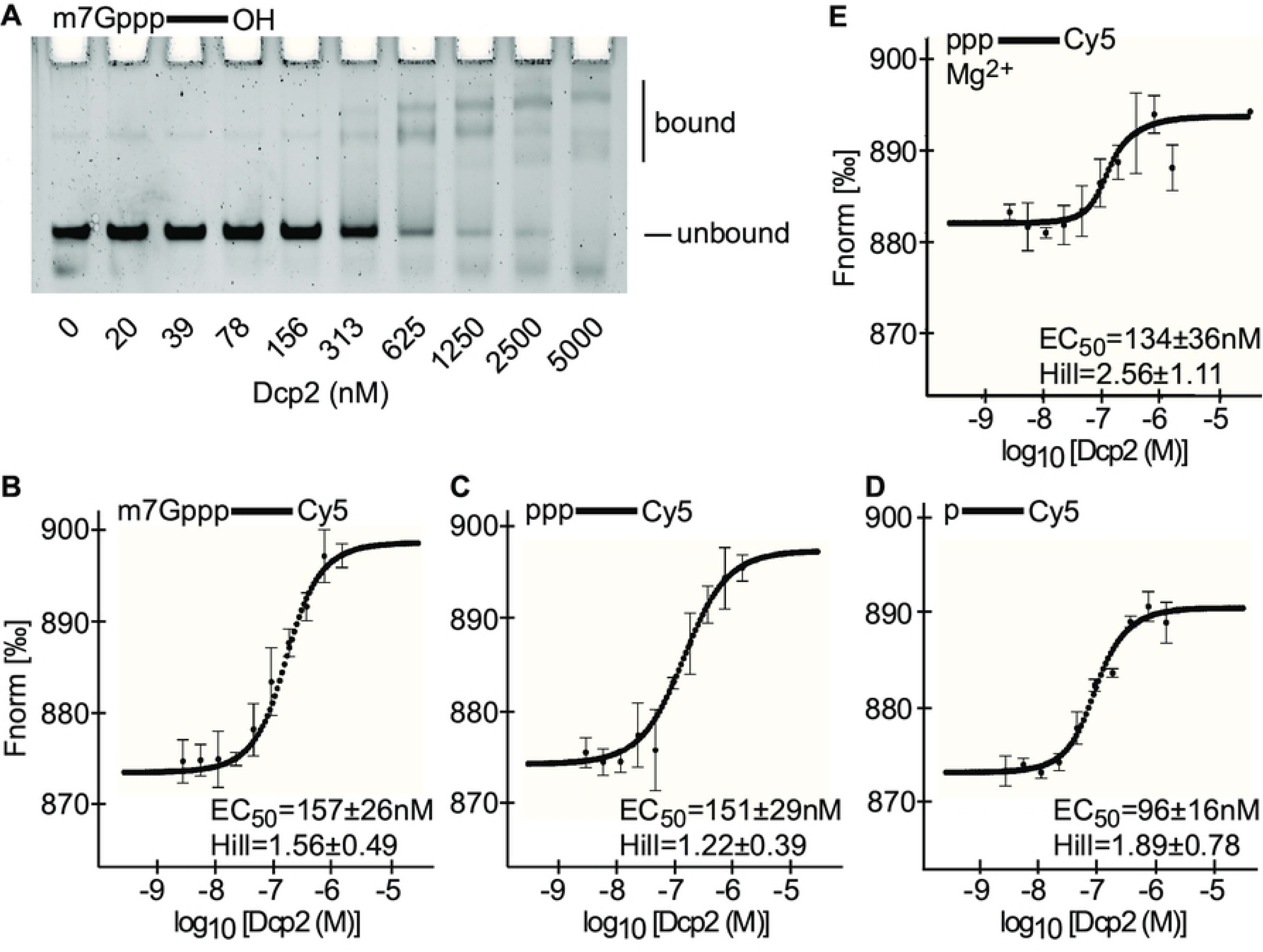
Dcp2 binds to RNA. A. Gel-shift assay. As indicated above the gel slice, m7G-capped 3’ unlabeled RNA oligos were incubated with the concentration of Dcp2 indicated under each lane. The bound and unbound fractions are specified by the bars to the right of the gel slice. B. MST assay using 3’ Cy5 labeled m7G-capped RNA. The Fnorm is specified on the y-axis, while the concentration of Dcp2 specified by the x-axis. Each circular point corresponds to the median Fnorm at that concentration of Dcp2, with the bars representing the s.d. for each data point, n=3 biological replicates. The median EC_50_ ± s.d. of the three replicates is indicated in the lower-right corner of the panel. The perforated line corresponds to the Hill model that was constructed based on the median of each datapoint. C. Same as panel A, except with 3’ Cy5 labeled 5’ triphosphate RNA. D. Same as panel A, except with 3’ Cy5 labeled 5’ monophosphate RNA. E. Same as panel C, except the binding buffer included Mg^2+^.

To determine the extent to which the m7G cap drives the affinity of Dcp2 for RNA, we repeated the MST experiment using oligos that were 5’ triphosphate. Here, we observe that Dcp2 exhibited a EC_50,obs_ of 151±29 nM for this RNA, with a Hill coefficient of 1.22±0.39 (Fig 2C). Next, we repeated this assay using RNA oligos that were 5’ monophosphate (Fig 2D). In this case, we determined that Dcp2 exhibited a EC_50,obs_ of 96±16 nM, with a Hill coefficient of 1.89±0.78. Thus, the affinity of Dcp2 for the reaction product (i.e. a 5’ monophosphate) is ∼1.6-fold stronger than the affinity of Dcp2 for the reaction substrate. To ensure that the lack of Mg^2+^ in the binding mixture did not majorly influence our results, we incubated Dcp2 with 5’ triphosphate RNA in our decapping buffer and repeated the MST assay, where we found that Mg^2+^ does not majorly influence the EC_50,obs_ of Dcp2 for this RNA (Figs 2C,E).

Multiple reports have found that mutating key residues (E147, E148) within the catalytic center of the human Dcp2 NUDIX domain abrogates catalytic decapping (29,34). It has been suggested that the NUDIX domain promotes RNA binding [39]. To test this, we purified human Dcp2^E147Q,E148Q^ and confirmed that this enzyme cannot de-cap RNA *in vitro* (S1A Fig, Fig 3A). We next repeated our MST assay, applying this nuclease deficient Dcp2 to m7G-capped RNA, where we observed no binding (Fig 3B).

**Figure 3.**
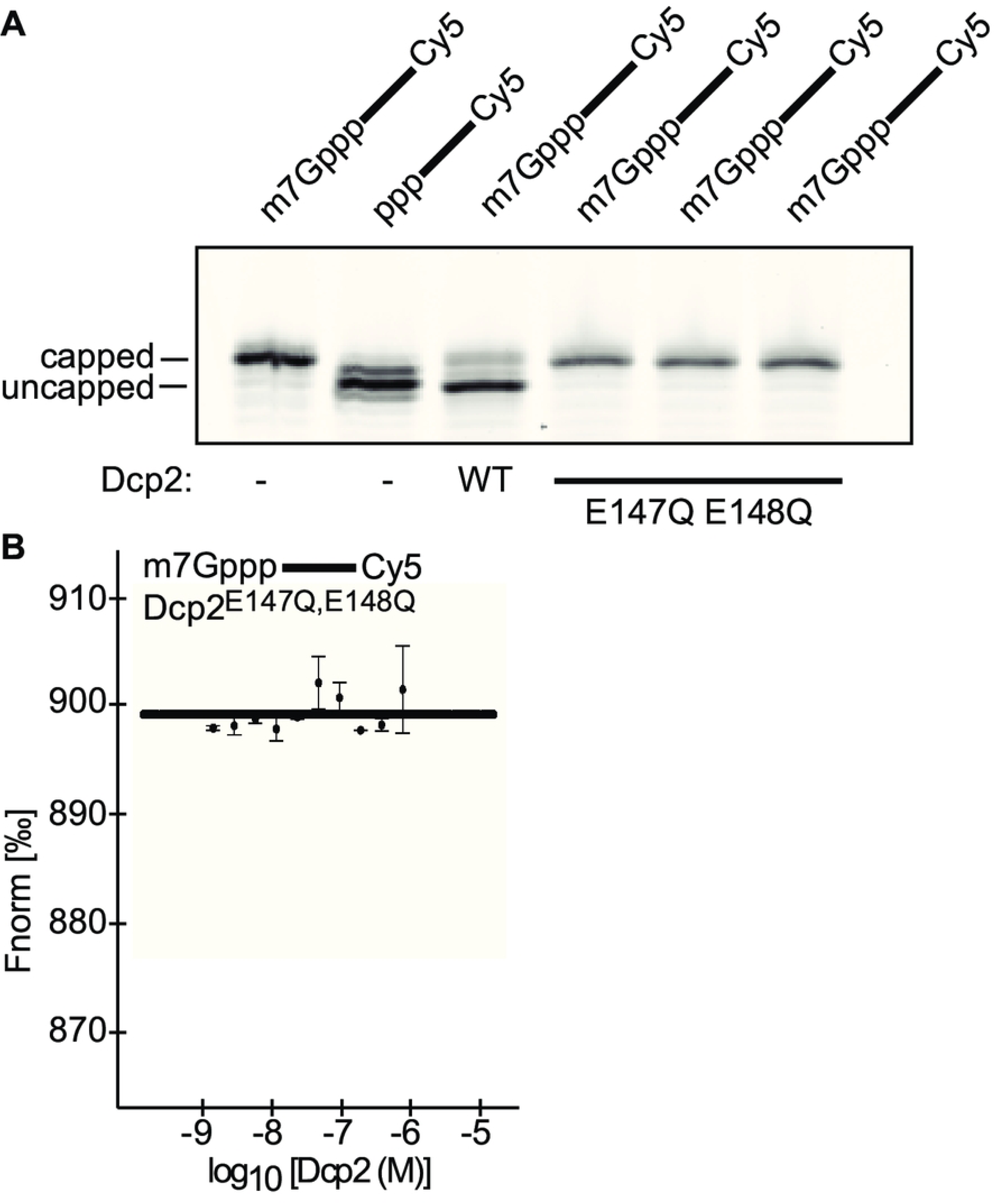
The NUDIX domain of Dcp2 participates in RNA binding. A. Decapping reactions using RNA oligos containing the 5’ moieties that are indicated above each lane. Experiments were performed in the absence (-), or presence of wild-type Dcp2, or mutant Dcp2, as specified under each lane. B. MST assay, with 3’ Cy5 labeled m7G-capped RNA. The Fnorm is specified on the y-axis, while the concentration of mutant Dcp2 specified by the x-axis. Each circular point corresponds to the median Fnorm at that concentration of Dcp2, with the bars representing the s.d. for each data point, n=2 biological replicates. The perforated line corresponds to the non-binder model that was constructed based on the median of each datapoint.

### Dcp2 binds to and de-caps m7G-capped RNA oligos that are bound to eIF4E

In the cytosol, the 5’ cap is associated with the translational initiation factor, eIF4E [2]. The contribution of bound eIF4E to mRNA decapping is unclear, although previous reports have found that eIF4E inhibits yeast decapping activity *in vitro* [10]. To test the effect of eIF4E on Dcp2-mediated decapping, we purified full-length human eIF4E to >90% purity (S2A Fig). Using a gel-shift assay, we confirmed that purified eIF4E binds specifically to m7G-capped RNA and does not interact with RNA lacking a cap structure, i.e. a 5’ triphosphate RNA, or with A-capped RNA (S2B Fig).

The effect of prebound eIF4E on Dcp2 RNA binding (see Fig 2) was then tested using a gel-shift assay. First, we incubated m7G-capped, 3’ unlabeled RNA with eIF4E, Dcp2, or no factor. We then resolved the bound and the unbound fractions using nondenaturing PAGE. Similar to the affinity experiments in Fig. 2, we conducted these experiments in the absence of divalent cation to prevent catalytic decapping (see Fig 1B). In these conditions, we observed a distinct eIF4E shift and multiple Dcp2 shifts (Fig 4A). We repeated these two experiments in the presence of excess m7GpppA cap analog, where we confirmed that cap analog completely inhibits the binding of eIF4E to RNA, and mildly inhibits the binding of Dcp2 to RNA (Fig 4A). Next, to test the effect of prebound eIF4E on Dcp2 binding, we pre-incubated the oligo with eIF4E, after which we added Dcp2 to the same concentration as eIF4E. Here, we observed gel-shifts corresponding to eIF4E bound singly to the oligo, as well as Dcp2 bound to the oligo (Fig 4A). Intriguingly, we noted that the amount of eIF4E bound singly to the oligo was decreased by ∼4.5-fold compared to the +eIF4E only condition. In addition, we noted that pre-bound eIF4E modulated the Dcp2 shift pattern. To better visualize these effects, we quantified the signal intensity across the +eIF4E, +Dcp2, and +eIF4E +Dcp2 lanes (Fig. 4B). We then repeated the +eIF4E +Dcp2 binding condition in the presence of excess m7GpppA, where we again confirmed that cap analog inhibits eIF4E (Fig 4A).

**Figure 4.**
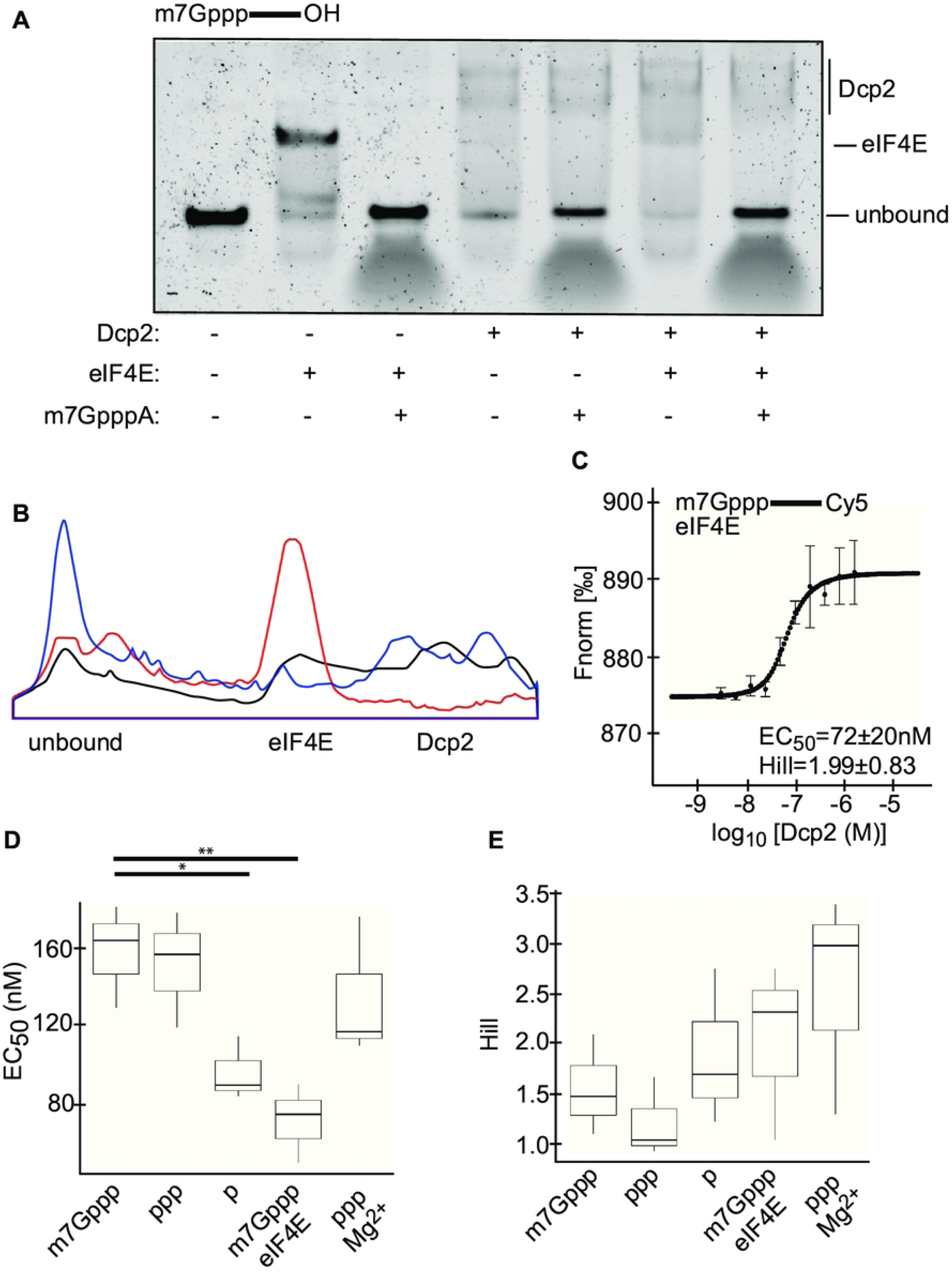
Dcp2 binds to RNA that is pre-bound to eIF4E. A. Gel-shift assays using m7G-capped 3’ unlabeled RNA oligos, as indicated above the gel slice. Experiments were performed in the absence or presence of Dcp2, eIF4E, and m7GpppA cap analog, as specified under each lane. The unbound fraction, the fraction bound singly to eIF4E, and the fractions bound to Dcp2 are specified by the bars to the right of the gel slice. B. Signal intensity was plotted across the +eIF4E lane (red), +Dcp2 lane (blue), and +eIF4E +Dcp2 lane (black) shown in panel A. C. MST assay, wherein m7G capped, 3’ Cy5 labeled RNA oligo and eIF4E, as indicated by the label in the upper-left corner, were incubated with a 2-fold serial dilution of Dcp2. The Fnorm is specified on the y-axis, while the concentration of Dcp2 specified by the x-axis. Each circular point corresponds to the median Fnorm at that concentration of Dcp2, with the bars representing the s.d. for each data point, n=3 biological replicates. The median EC_50_ ± s.d. of the three replicates is indicated in the lower-right corner of the panel. The perforated line corresponds to the Hill model that was constructed based on the median of each datapoint. D. Box plot of the EC_50_ ± s.d. of Dcp2 for each of the substrates. Box, first to last quartiles; whiskers, 1.5× interquartile range; centre line, median; points, outliers. Unpaired Wilcoxon rank sum tests were conducted, with results by the stars above the two horizontal black lines. E. Same as panel D, except showing the Hill coefficient ± s.d. of Dcp2 for each of the substrates.

To more quantitatively assay the effect of bound eIF4E on the affinity of Dcp2 for RNA, we pre-incubated m7G-capped, 3’ Cy5 labeled RNA with eIF4E, after which we incubated this mixture with a serial dilution of Dcp2. The affinity of Dcp2 for the RNA was then measured using MST, where we found the EC_50,obs_ between Dcp2 and the m7G-capped RNA was 72±20 nM in the presence of eIF4E, with a Hill coefficient of 1.99±0.83 (Fig 4C). Thus, eIF4E increases the affinity of Dcp2 for the RNA oligo by ∼2-fold (Fig 3).

Next, we tested the influence of prebound eIF4E on Dcp2 decapping rate under single-hit kinetics (k_obs_). When determining k_obs_, it is necessary to hold the concentration of Dcp2 at a level that is saturating (39). To determine at which point Dcp2 becomes saturating, we incubated the m7G-capped RNA oligos with increasing concentrations of Dcp2 and an excess of Xrn1 (S3 Fig). Through this experiment, we found that Dcp2 was saturating at concentrations at least 10-fold in excess of the oligo. Thus, we held Dcp2 at a 20-fold molar excess to quantitate the k_obs_. Using this molar ratio, we conducted a decapping time-course assay, wherein we incubated Dcp2 with the m7G-capped RNA oligo for various time-points before stopping the reaction. Upon the completion of the time-course, we separated the capped from the uncapped RNA species via denaturing PAGE (Fig 5A). By comparing the ratio of the capped to the uncapped species across the time-course, we found that the k_obs_ of human Dcp2 is 0.05±0.01^min^ (Fig 4B). This rate is ∼5X higher than the k_obs_ of the yeast Dcp1:Dcp2 heterodimer [14]. To quantify the effect of eIF4E on the rate of decapping by Dcp2, we first incubated the m7G-capped RNA oligo with a 20-fold molar excess of eIF4E, before adding Dcp2 and incubating the mixture across the same time-course as was used above (Fig 5B, C). From this experiment, we calculated the k_obs_ of Dcp2 on eIF4E bound RNA as being 0.03±0.005^min^. Thus, eIF4E does not significantly impede, nor promote, the catalytic activity of Dcp2 in vitro.

**Figure 5.**
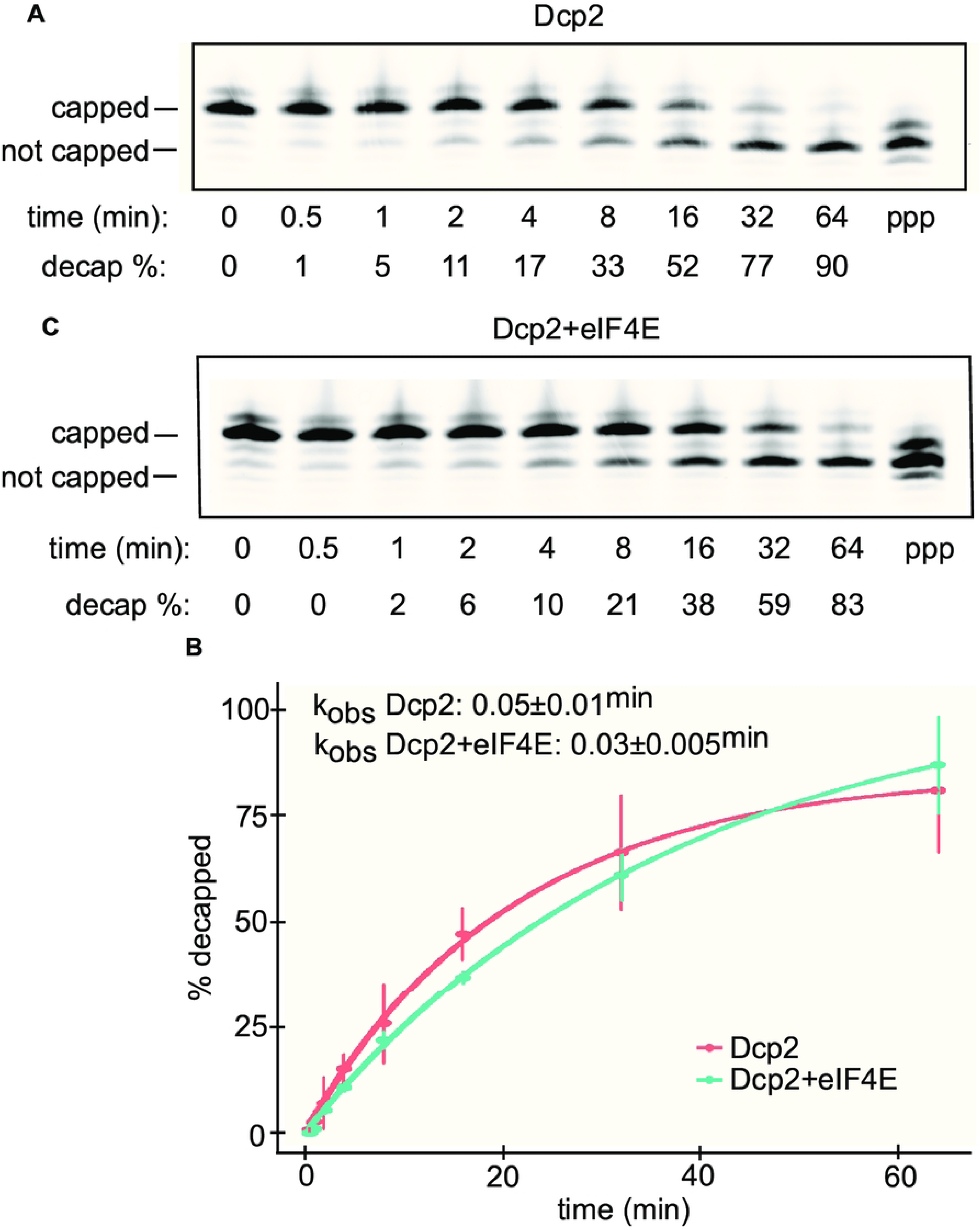
Dcp2 de-caps RNA oligos that are bound to eIF4E. A. Decapping time-course reactions using m7G-capped RNA oligos. RNA oligo was incubated with Dcp2 for the amount of time specified under each lane. The decapping % for the data is shown underneath each lane. The capped and not-capped RNA species are indicated by the horizontal bars to the right of the gel slice. The right-most lane contains 5’ triphosphate RNA oligo. B. Point-and-line plot of the decapping time-course data. Each circular point corresponds to the median % decapped after that amount of time with Dcp2, with the bars representing the s.d. for each data point, n=3 biological replicates. The red data corresponds to the data collected using Dcp2, while the blue data corresponds to the data collected using both eIF4E and Dcp2. The median k_obs_ ± s.d. of the three replicates is indicated in the top left corner of the plot. C. Same as panel A, except showing the RNA oligo incubated with eIF4E prior to Dcp2.

## DISCUSSION

Our results provide an additional level of mechanistic detail on the activity of Dcp2 in humans. We confirmed that this enzyme specifically removes the m7G cap from RNA in the absence of any decapping coactivators, with a turnover rate of ∼20 minutes under single-order kinetics (Fig 3A). Several recent studies have reported that human Dcp2 decaps RNA in the absence of any decapping coactivators in vitro, although the extent of this activity is conflicting across various reports [22,28]. These distinctions can likely be attributed to differences in the employed experimental methodology. How additional human decapping coactivators impact the rate of decapping remains to be determined.

Surprisingly, we found that Dcp2 has an affinity for m7G-capped RNA (Figs 2A, B, 4D) that is in-line with the affinity of eIF4E for m7G-capped RNA [29,31]. In addition, we found that Dcp2 has an approximately equal affinity for 5’ triphosphate RNA (Figs 2C, 4D). These latter results are consistent with previous reports, which have found that human Dcp2 binds to both capped and 5’ triphosphate RNA, and that 5’ triphosphate RNA competes with capped RNA for Dcp2 binding [27,32]. Intriguingly, we found that Dcp2 has a slightly stronger affinity for 5’ monophosphate RNA (Figs 2D, 4D). Structural studies of yeast Dcp2 have found that this enzyme contains a bipartite active site and forms distinct ‘on’ and ‘off’ states, with only the ‘on’ state capable of catalyzing decapping [19,20,41]. How these different structures interact with these binding options will require further examination. Moreover, our results suggest that Dcp2 binds to RNA cooperatively and/or nonspecifically in all of the tested conditions (see Hill coefficients in Figs 2, 4), which is in-line with previous findings obtained using yeast Dcp2 [39]. In addition, our results imply that Dcp2 requires a functional NUDIX domain to bind RNA (Fig. 3), which has also been suggested by others [39]. It should be noted that Mg^2+^ is widely understood to influence protein-RNA interactions. While we found that Mg^2+^ does not majorly influence the binding of Dcp2 (Fig 2C, E), nor eIF4E (Fig 4A, S3B Fig) to RNA, these experiments cannot exclude the possibility that Mg^2+^ will influence how Dcp2 and eIF4E interact on an RNA scaffold.

Perhaps most surprising, we found that bound eIF4E increases the affinity of Dcp2 for the RNA by ∼2-fold (Fig 4C). It is currently unknown how Dcp2 is recruited to the mRNA 5’ ends. While our data is limited to in vitro examination, our results indicate that eIF4E may play a role in this process. This makes intuitive sense, as eIF4E serves as the 5’ end beacon for the translation initiation apparatus. Our results suggest that Dcp2 binding results in eIF4E dissociation (Fig 4A). This model is consistent with both our finding that pre-bound eIF4E does not significantly impact the rate of Dcp2 decapping (Fig 5), as well as the numerous studies, which suggest that Dcp2 cannot access the cap when eIF4E is bound [10,19–21,42–46]. How exactly these two factors interact with each other on the same RNA remains to be determined.

## EXPERIMENTAL PROCEDURES

Full experimental procedures can be found in the supporting information

## DATA AVAILABILITY

All data are contained in the article.

## ACKNOWLEDGEMENTS

We would like to thank Dr. Jordana Henderson and her team at TriLink Biotechnologies for their help in making the RNA oligos used in this study. We would like to thank Dr. Megerditch Kiledjian for his help in establishing a protocol to obtain highly pure Dcp2.

## AUTHOR CONTRIBUTIONS

ZM designed the experiments, conducted the experiments, analyzed the data, wrote the manuscript, and edited the manuscript. JC supervised the project, acquired funding, and edited the manuscript.

## REFERENCES

1. Ramanathan A., Robb GB., Chan SH. (2016) mRNA capping: biological functions and applications. Nucleic Acids Res. 2016;44(16):7511-26.

2. Jackson RJ, Hellen CUT, Pestova TV. The mechanism of eukaryotic translation initiation and principles of its regulation. Nat Rev Mol Cell Biol. 2010;11(2):113–127.

3. Nagarajan VK, Jones CI, Newbury SF, Green PJ. XRN 5′→3′ exoribonucleases: Structure, mechanisms and functions. Biochim Biophys Acta. 2013;1829(0):590–603.

4. Tuck AC, Rankova A, Arpat AB., Liechti, L. A., Hess, D., Iesmantavicius, V., et al. Mammalian RNA Decay Pathways Are Highly Specialized and Widely Linked to Translation. Mol Cell. 2020;77(6):1222–1236.e13.

5. Bae H, Coller J. Codon optimality-mediated mRNA degradation: Linking translational elongation to mRNA stability. Mol Cell. 2022;82(8):1467–1476.

6. Pelechano V, Wei W, Steinmetz LM. Widespread Co-translational RNA Decay Reveals Ribosome Dynamics. Cell. 2015;161(6):1400–12.

7. Hu W, Sweet TJ, Chamnongpol S, Baker KE, Coller J. Co-translational mRNA decay in *Saccharomyces cerevisiae*. Nature. 2009;461(7261):225-9.

8. D’Orazio KN, Wu CCC, Sinha N, Loll-Krippleber R, Brown GW, Green R. The endonuclease Cue2 cleaves mRNAs at stalled ribosomes during No Go Decay. Elife. 2019;8:e49117.

9. Hu W, Petzold C, Coller J, Baker KE. Nonsense-mediated mRNA decapping occurs on polyribosomes in *Saccharomyces cerevisiae*. Nat Struct Mol Biol. 2017;17(2):244–7.

10. Schwartz DC, Parker R. mRNA Decapping in Yeast Requires Dissociation of the Cap Binding Protein, Eukaryotic Translation Initiation Factor 4E. Mol Cell Biol. 2000;20(21):7933-42.

11. Absmeier E, Chandrasekaran V, O’Reilly FJ, Stowell JAW, Rappsilber J, Passmore LA (2022) Specific recognition and ubiquitination of slow-moving ribosomes by human CCR4-NOT. Nat Struct Mol Biol. 2023;30:1314–1322.

12. Braun JE, Truffault V, Boland A, Huntzinger E, Chang CT, Haas G, et al. A direct interaction between DCP1 and XRN1 couples mRNA decapping to 5′ exonucleolytic degradation. Nat Struct Mol Biol. 2012;19(12):1324–31.

13. Wurm JP, Sprangers R. Dcp2: an mRNA decapping enzyme that adopts many different shapes and forms. Curr Opin Struct Biol. 2019 Dec;59:115–123.

14. Paquette DR, Tibble RW, Daifuku TS, Gross JD. Control of mRNA decapping by autoinhibition. Nucleic Acids Res. 2018;46(12):6318–6329.

15. Lobel JH, Tibble RW, Gross JD. Pat1 activates late steps in mRNA decay by multiple mechanisms. Proc Natl Acad Sci U S A. 2019;116(47):23512–23517.

16. Charenton C, Gaudon-Plesse C, Fourati Z, Taverniti V, Back R, Kolesnikova O, et al. A unique surface on Pat1 C-terminal domain directly interacts with Dcp2 decapping enzyme and Xrn1 5′–3′ mRNA exonuclease in yeast. Proc Natl Acad Sci U S A. 2017;114(45):E9493–E9501.

17. He F, Jacobson A. Control of mRNA decapping by positive and negative regulatory elements in the Dcp2 C-terminal domain. RNA. 2015;21(9):1633–47.

18. Vidya E, Duchaine TF. Eukaryotic mRNA Decapping Activation. Front Genet. 2022;13:832547.

19. Valkov E, Muthukumar S, Chang CT, Jonas S, Weichenrieder O, Izaurralde E. Structure of the Dcp2–Dcp1 mRNA-decapping complex in the activated conformation. Nat Struct Mol Biol. 2016;23(6):574–9.

20. Mugridge JS, Tibble RW, Ziemniak M, Jemielity J, Gross JD. Structure of the activated Edc1-Dcp1-Dcp2-Edc3 mRNA decapping complex with substrate analog poised for catalysis. Nat Commun. 2018;9(1):1152.

21. Charenton C, Taverniti V, Gaudon-Plesse C, Back R, Séraphin B, Graille M. Structure of the active form of Dcp1–Dcp2 decapping enzyme bound to m7GDP and its Edc3 activator. Nat Struct Mol Biol. 2016;23(11):982–986.

22. Dunckley T. The DCP2 protein is required for mRNA decapping in *Saccharomyces cerevisiae* and contains a functional MutT motif. EMBO J. 1999;18(19):5411–22.

23. Chang CT, Bercovich N, Loh B, Jonas S, Izaurralde E. The activation of the decapping enzyme DCP2 by DCP1 occurs on the EDC4 scaffold and involves a conserved loop in DCP1. Nucleic Acids Res. 2014;42(8):5217–33.

24. Fenger-Grøn M, Fillman C, Norrild B, Lykke-Andersen J. Multiple Processing Body Factors and the ARE Binding Protein TTP Activate mRNA Decapping. Mol Cell. 2005;20(6):905–15.

25. Yu JH, Yang WH, Gulick T, Bloch KD, Bloch DB, Yu JH, et al. Ge-1 is a central component of the mammalian cytoplasmic mRNA processing body. RNA. 2005;11(12):1795–802.

26. Li Y, Ho ES, Gunderson SI, Kiledjian M. Mutational analysis of a Dcp2-binding element reveals general enhancement of decapping by 5′-end stem-loop structures. Nucleic Acids Res. 2009;37(7):2227–37.

27. Li Y, Song MG, Kiledjian M. Transcript-Specific Decapping and Regulated Stability by the Human Dcp2 Decapping Protein. Mol Cell Biol. 2008;28(3):939–48.

28. Wang Z, Jiao X, Carr-Schmid A, Kiledjian M. The hDcp2 protein is a mammalian mRNA decapping enzyme. Proc Natl Acad Sci U S A. 2002;99(20):12663–12668.

29. Drazkowska K, Tomecki R, Warminski M, Baran N, Cysewski D, Depaix A, et al. 2′-*O*-Methylation of the second transcribed nucleotide within the mRNA 5′ cap impacts the protein production level in a cell-specific manner and contributes to RNA immune evasion. Nucleic Acids Res. 2022;50(16):9051–9071.

30. Tamarkin-Ben-Harush A, Vasseur J, Debart F, Ulitsky I, and Dikstein R. Cap-proximal nucleotides via differential eIF4E binding and alternative promoter usage mediate translational response to energy stress. eLife. 2017;6:e21907.

31. van Dijk E. Human Dcp2: a catalytically active mRNA decapping enzyme located in specific cytoplasmic structures. EMBO J. 2002;21(24):6915–24.

32. Piccirillo C, Khanna R, Kiledjian M. Functional characterization of the mammalian mRNA decapping enzyme hDcp2. RNA. 2003;9(9):1138–1147.

33. Mauer J, Luo X, Blanjoie A, Jiao X, Grozhik AV, Patil DP, et al. Reversible methylation of m6Am in the 5′ cap controls mRNA stability. Nature. 2017;541(7637):371-375.

34. Huang L, Zhang C. Microscale Thermophoresis (MST) to Detect the Interaction Between Purified Protein and Small Molecule. Methods Mol Biol. 2021;2213:187–193.

35. Weiss JN. The Hill equation revisited: uses and misuses. FASEB J. 1997;11(11):835–41.

36. Davidovich C, Wang X, Cifuentes-Rojas C, Goodrich KJ, Gooding AR, Lee JT, et al. Toward a consensus on the binding specificity and promiscuity of PRC2 for RNA. Mol Cell. 2015;57(3):552–8.

37. Jarmoskaite I, AlSadhan I, Vaidyanathan PP, Herschlag D. How to measure and evaluate binding affinities. eLife. 2020;9:e57264.

38. Deshmukh MV, Jones BN, Quang-Dang D, Flinders J, Floor SN, Kim C, et al. mRNA decapping is promoted by an RNA-binding channel in Dcp2. Mol Cell. 2008;29(3):324–36.

39. Jones BN, Quang-Dang DU, Oku Y, Gross JD. Chapter 2: A Kinetic Assay to Monitor RNA Decapping Under Single-Turnover Conditions. Methods Enzymol. 2008:448:23–40.

40. Charenton C, Taverniti V, Gaudon-Plesse C, Back R, Séraphin B, Graille M. Structure of the active form of Dcp1–Dcp2 decapping enzyme bound to m7GDP and its Edc3 activator. Nat Struct Mol Biol. 2016;23(11):982–986.

41. Grüner S, Peter D, Weber R, Wohlbold L, Chung MY, Weichenrieder O, et al. The Structures of eIF4E-eIF4G Complexes Reveal an Extended Interface to Regulate Translation Initiation. Mol Cell. 2016;64(3):467–479.

42. Kinkelin K, Veith K, Grünwald M, Bono, F. Crystal structure of a minimal eIF4E–Cup complex reveals a general mechanism of eIF4E regulation in translational repression. RNA. 2012;18(9):1624–1634

43. Marcotrigiano J, Gingras AC, Sonenberg N, Burley SK. Cocrystal Structure of the Messenger RNA 5′ Cap-Binding Protein (eIF4E) Bound to 7-methyl-GDP. Cell. 1997;89(6):951–61.

44. Papadopoulos E, Jenni S, Kabha E, Takrouri KJ, Yi T, Salvi N, et al. Structure of the eukaryotic translation initiation factor eIF4E in complex with 4EGI-1 reveals an allosteric mechanism for dissociating eIF4G. Proc Natl Acad Sci U S A. 2014;111(31):E3187–95.

45. Tomoo K, Shen X, Okabe N, Nozoe Y, Fukuhara S, Morino S, et al. Crystal structures of 7-methylguanosine 5′-triphosphate (m7GTP)- and P1-7-methylguanosine-P3-adenosine-5′,5′-triphosphate (m7GpppA)-bound human full-length eukaryotic initiation factor 4E: biological importance of the C-terminal flexible region. Biochem J. 2002;362(Pt 3):539-44.

46. Yakhnin AV, Yakhnin H, Babitzke P. Gel Mobility Shift Assays to Detect Protein–RNA Interactions. Methods Mol Biol. 2012:905:201–11.

